# Metal-binding propensity in the mitochondrial dynamin-related protein 1

**DOI:** 10.1101/2022.01.15.476484

**Authors:** Krishnendu Roy, Thomas J. Pucadyil

## Abstract

Dynamin-related protein1 (Drp1) functions to divide mitochondria and peroxisomes by binding specific adaptor proteins and lipids, both of which are integral to the limiting organellar membrane. In efforts to understand how such multivalent interactions regulate Drp1 functions, *in vitro* reconstitution schemes rely on recruiting soluble portions of the adaptors appended with genetically encoded polyhistidine tags onto membranes containing Ni^2+^-bound chelator lipids. These strategies are facile and circumvent the challenge in working with membrane proteins but assume that binding is specific to proteins carrying the polyhistidine tag. Here, we find using chelator lipids and chelator beads that both native and recombinant Drp1 directly bind Ni^2+^ ions. Metal-binding therefore represents a potential strategy to deplete or purify Drp1 from native tissue lysates. Importantly, high concentrations of the metal in solution inhibit GTP hydrolysis and renders Drp1 inactive in membrane fission. Together, our results emphasize a metal-binding propensity, which could significantly impact Drp1 functions.

## Introduction

The dynamin-related protein 1 (Drp1) functions to affect mitochondrial and peroxisomal division. Drp1 is a soluble protein that is recruited by specific adaptors such as Mitochondrial fission factor (Mff) and the Mitochondrial dynamics protein of 49 kDa (Mid49) and 51 kDa (Mid51), which are single-pass integral membrane proteins present on the outer mitochondrial membrane (Kraus et al. 2021). Drp1 interacts with these adaptor proteins through conserved interfaces on the G and stalk domains as well as with membrane lipids through an approximately 100-residue-long unstructured region known as the variable domain (Ramachandran 2018; Kalia and Frost 2019; Kraus et al. 2021). Soluble members of the dynamin superfamily bind membranes and self-assemble into helical scaffolds with enhanced GTPase activity (Chappie and Dyda 2013). GTPase-induced conformational changes in the scaffold forces membrane constriction leading to fission (Dar et al. 2015).

Previous reconstitution approaches have relied on membrane templates to understand Drp1 functions. The simplest of these rely on lipid-based recruitment of Drp1 on to the membrane and have demonstrated Drp1’s capacity for constricting and severing membrane tubes (Kamerkar et al. 2018a). In attempts to involve adaptor proteins in this scheme, but circumvent experimental challenges in working with integral membrane proteins, our group and several others have used a strategy to recruit polyhistidine (6xHis) tagged constructs of the soluble domains of the adaptors onto templates containing the Ni^2+^-bound chelator lipid 1,2-dioleoyl-*sn*-glycero-3-[(N-(5-amino-1-carboxypentyl)iminodiacetic acid)succinyl] (nickel salt) (DGS-NTA(Ni^2+^)) (Koirala et al. 2013; Clinton et al. 2016; Osellame et al. 2016; Clinton and Mears 2017; Kamerkar et al. 2018a). However, a previous study utilizing such a strategy reported a direct interaction between Drp1 and DGS-NTA(Ni^2+^), which was surprising because the Drp1 construct used was devoid of a 6xHis tag (Koirala et al. 2013). Such interactions could potentially complicate efforts to reconstitute Drp1 functions and we therefore probed its molecular basis.

## Materials and Methods

### Protein Expression and Purification

Human Drp1 (Uniprot ID: O00429-4) with a C-terminal StrepII tag in pET15b (Addgene plasmid #174428) was expressed in BL21(DE3) and grown in autoinduction medium at 18°C for 36 hours. Bacterial cells were pelleted and stored at −40 °C. The frozen bacterial pellet was thawed in 20 mM HEPES pH 7.4, 500 mM NaCl with a protease inhibitor cocktail tablet (Roche) and lysed by sonication in an ice-water bath. Lysates were spun at 30,000x g for 20 mins. The supernatant was applied to a 5 ml streptactin column (GE Healthcare), pre-equilibrated with the same buffer and washed extensively. Buffer was exchanged to 20mM HEPES pH 7.4, 150mM NaCl and bound protein was eluted with the same buffer containing 2.5 mM desthiobiotin (Sigma-Aldrich). Proteins were spun at 100,000x g to remove aggregates before use in assays. Protein concentration was estimated from UV absorbance at 280 nm from the molar extinction coefficient predicted using the Expasy ProtParam tool.

### Liposome Preparation

1,2-dioleoyl-*sn*-glycero-3-phosphocholine (DOPC), 1’,3’-bis[1,2-dioleoyl-*sn*-glycero-3-phospho]-glycerol (18:1 cardiolipin, CL) and 1,2-dioleoyl-*sn*-glycero-3-[(N-(5-amino-1-carboxypentyl)iminodiacetic acid)succinyl] (nickel salt) (DGS-NTA(Ni^2+^) were purchased from Avanti Polar Lipids. DOPC was the bulk lipid in all experiments. The UV-activable, diazirine-containing fluorescent lipid probe (BODIPY-diazirine PE) was prepared according to published protocols (Jose and Pucadyil 2020; Jose et al. 2020). Desired lipids were aliquoted in the required amount into a glass tube and dried under high vacuum for 30 mins to a thin film. Dried lipids were then hydrated in deionized water at 50 °C for 30 mins, vortexed vigorously and extruded through 100 nm pore-size polycarbonate filters (Whatman).

### Proximity-based Labeling of Membrane-Associated Proteins (PLiMAP)

Typical experiments involved mixing liposomes containing BODIPY-diazirine PE (1 mol%) with proteins at a 100:1 molar ratio in a final volume of 30 μl. The reaction was incubated in dark at room temperature for 30 mins. For experiments with 6xHis-mCherry, Drp1 was added to liposomes after they were incubated with the 6xHis-mCherry for 30 mins. The reaction mix was exposed to 365 nm UV light (UVP crosslinker CL-1000L) at an intensity of 200 mJ.cm^-2^ for 1 min. The reaction mix was then boiled with sample buffer and resolved using SDS-PAGE. Gels were first imaged for BODIPY fluorescence and later fixed and imaged for Coomassie Brilliant Blue stain on an iBright 1500 (Thermo Fisher).

### GTPase Assays

Drp1 (1 μM) was mixed with GTP (1 mM) and MgCl_2_ (1 mM) in 20 mM HEPES pH 7.4, 150 mM NaCl in the absence or presence of liposomes and incubated at 37 °C. Buffers contained increasing concentrations of NiCl_2_ to test the effect of Ni^2+^ on Drp1’s GTPase activity. Aliquots were taken out at regular intervals, quenched with EDTA and the released inorganic phosphate was assayed with a malachite-green reagent (Leonard et al. 2005).

### Supported Membrane Nanotubes (SMrT) and Fluorescence Imaging

SMrT templates were prepared as described previously (Dar et al. 2017). Briefly, lipids were aliquoted at the desired ratios and brought to a final concentration of 1 mM in chloroform. Lipid mixes contained a small amount (1 mol%) of the fluorescent lipid probe *p*Texas-Red DHPE (Thermo Fisher). 2 μl of the lipid mix was spread with a glass syringe on a PEGylated glass coverslip, dried and assembled inside an FCS2 flow chamber (Bioptechs). The flow cell was then filled with buffer to hydrate the dried lipid. SMrT templates were formed by extruding the hydrated membrane into narrow tubes by flowing buffer at high flow rates. Fluorescence images were recorded before and after flowing Drp1 (1 μM) with GTP (1 mM) and MgCl2 (1 mM) in 20 mM HEPES pH 7.4, 150 mM NaCl. Buffers contained NiCl2 (1 mM) to test the effect of Ni^2+^ on Drp1’s fission activity. Imaging was carried out through 100x, 1.4 NA oil-immersion objective on an Olympus IX83 inverted microscope attached with a stable LED light source (CoolLED) and an Evolve 512 EMCCD camera (Photometrics). Image acquisition was controlled by Micro-Manager and rendered using Fiji (Schindelin et al. 2012).

### Preparation of Brain Lysates

Lysates from adult goat brains were prepared as described earlier (Wu et al. 2010; Kamerkar et al. 2018b). Briefly, brain tissues were cleaned of meninges and blood vessels and homogenized in a Waring blender in breaking buffer (25 mM Tris pH 8.0, 500 mM KCl, 250 mM sucrose, 2 mM EGTA, and 1 mM DTT) with a protease inhibitor cocktail tablet (Roche). The lysate was spun at 160,000x g for 2 h. The supernatant was collected and desalted on a G-50 column into 20 mM HEPES pH 7.4, 150 mM KCl. Lysates were flash-frozen with 10% glycerol in liquid nitrogen and stored at −80 °C. Total protein content was estimated using the Pierce BCA Protein Assay Kit (Thermo Fisher).

### Pulldown with Chelator Beads

Co^2+^-bound NTA-sepharose beads (Takara) were stripped off the metal ion with 100 mM EDTA and used as NTA beads or charged with 100 mM NiCl_2_ and used as NTA(Ni^2+^) beads. Beads were equilibrated with 20 mM HEPES pH 7.4, 150 mM KCl and incubated with 0.4 ml of brain lysate (0.5 mg/ml) for 30 mins at room temperature. Samples were spun at 3000x g for 1 min and 20 μl of the supernatant was removed. Beads were then washed 3 times with 0.4 ml of the same buffer. The supernatant and beads were mixed with sample buffer, resolved using SDS-PAGE and probed for Drp1 using Western blotting.

### Western Blotting

Proteins were transferred onto PVDF membrane and blocked with 5% skimmed milk in TBST for 1 hr at room temperature. Anti-Drp1 antibody (3B5, Abcam, cat. no.: Ab56788) was used at a dilution recommended by the manufacturer in blocking buffer. The membrane was incubated with the primary antibody in blocking buffer for 3 hrs at room temperature. Subsequently, the membrane was washed with TBST and incubated with HRP-conjugated secondary antibody, diluted in blocking buffer for 1 hr at room temperature. Blots were rinsed with TBST and developed with the chemiluminescent substrate (WestPico, Thermo Fisher Scientific) and imaged on an iBright 1500 (Thermo Fisher).

## Results

The previously reported interaction between Drp1 and DGS-NTA(Ni^2+^) was assayed using liposome sedimentation (Koirala et al. 2013). To confirm this interaction using sensitive methods, we turned to a recently developed crosslinking approach called Proximity-based Labeling of Membrane-Associated Proteins (PLiMAP) (Jose and Pucadyil 2020; Jose et al. 2020). PLiMAP utilizes liposomes containing a small fraction (1 mol%) of a UV-activable, diazirine-containing fluorescent lipid probe (BODIPY-diazirine PE). Liposome binding brings proteins in proximity to the fluorescent probe and a brief exposure to UV covalently crosslinks the protein with the fluorescent probe. Liposome binding can then be quantified by in-gel fluorescence of the crosslinked protein resolved using SDS-PAGE.

We incubated recombinant Drp1 (1 μM) with liposomes (100 μM) containing increasing concentrations of DGS-NTA(Ni^2+^) and subjected them to PLiMAP. As shown in Fig. 1A (and quantitated in Fig. 1B), Drp1 is rendered fluorescent in the presence of liposomes containing DGS-NTA(Ni^2+^), but not in its absence. Moreover, Drp1 fluorescence increases with higher concentrations of DGS-NTA(Ni^2+^), implying that Drp1 binds DGS-NTA(Ni^2+^). Quantitation of Drp1 fluorescence with increasing concentration of liposomes containing 10 mol% DGS-NTA(Ni^2+^) revealed an apparent binding affinity of ~3 μM with a hill coefficient of 1.3 (Fig. 1C, D).

**Fig. 1.**
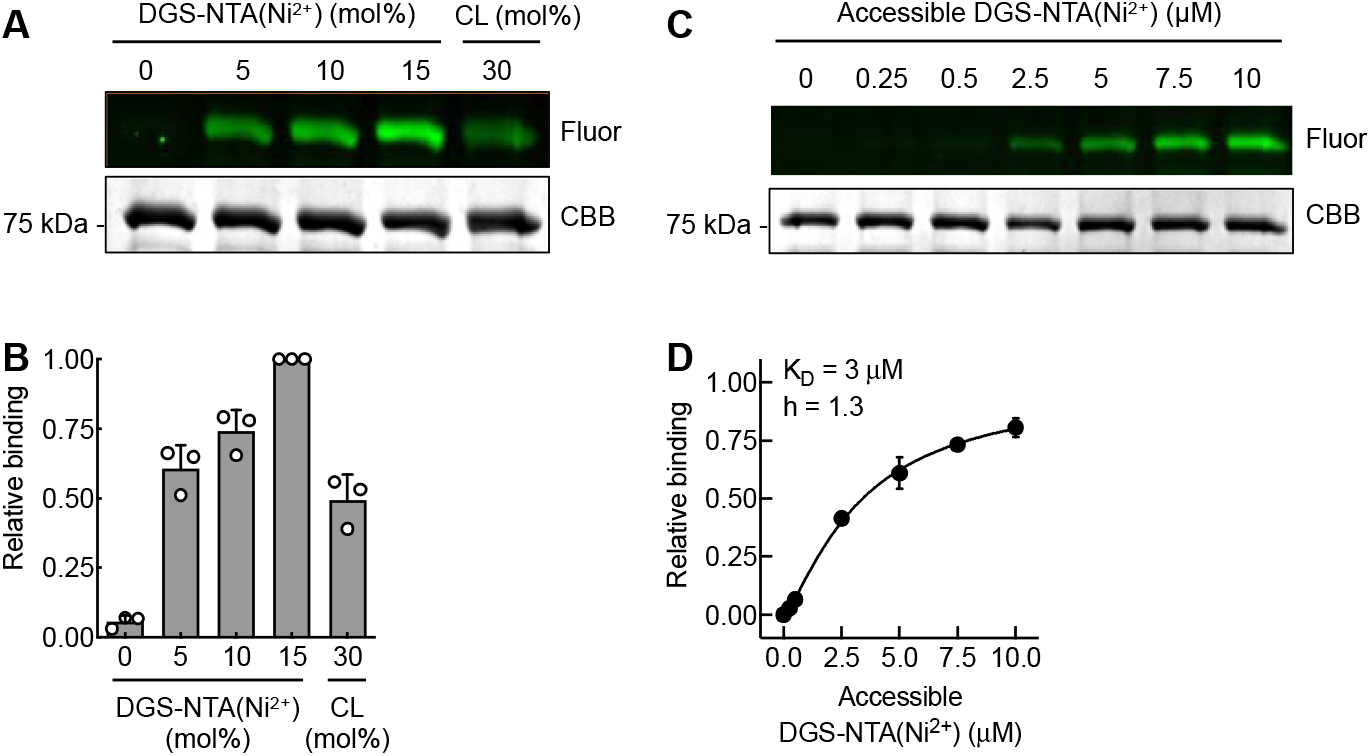
Drp1 binds the chelator lipid DGS-NTA(Ni^2+^). (A) Results from a representative PLiMAP experiment showing in-gel fluorescence (Fluor) and Coomassie brilliant blue (CBB) staining of Drp1 incubated with liposomes containing the indicated concentrations of DGS-NTA(Ni^2+^) and cardiolipin (CL). (B) Quantitation of in-gel fluorescence, which equates to Drp1 binding to liposomes. Data is normalized to binding seen with 15 mol% of DGS-NTA(Ni^2+^) and represents the mean ± SD of three independent experiments. (C) Results from a representative PLiMAP experiment showing in-gel fluorescence (Fluor) and Coomassie brilliant blue (CBB) staining of Drp1 incubated with increasing concentrations of 10 mol% DGS-NTA(Ni^2+^)-containing liposomes. (D) Fraction of bound Drp1 plotted against the accessible DGS-NTA(Ni^2+^) and fitted to a one-site specific binding with a hill slope equation. Data represents the mean ± SD of three independent experiments and is normalized to the fitted maximal binding (B_max_).

Drp1 functions have been assayed *in vitro* based on its ability to bind the mitochondrial lipid cardiolipin (CL) (Macdonald et al. 2014; Bustillo-Zabalbeitia et al. 2014; Stepanyants et al. 2015; Francy et al. 2017; Kamerkar et al. 2018a; Mahajan et al. 2021). Remarkably, we find that liposomes containing 5 mol% of DGS-NTA(Ni^2+^) recruited as much Drp1 as did liposomes containing 30 mol% CL (Fig. 1A, B), implying that Drp1 has higher affinity for DGS-NTA(Ni^2+^) than CL. Soluble dynamin proteins bind membranes and self-assemble into helical scaffolds, which in turn reorients catalytic residues in their GTP-binding G domain and stimulates their basal GTPase activity (Chappie and Dyda 2013). Thus, a stimulation in GTPase activity with membranes represents a read-out for dynamin self-assembly. Consistent with previous findings reporting that Drp1 organizes into helical scaffolds on CL-containing membranes (Francy et al. 2017), we find a 15-fold stimulation in Drp1’s GTPase activity in presence of 30% CL-containing liposomes (Fig. 2A). Despite its strong binding to DGS-NTA (Ni^2+^), Drp1’s GTPase activity only showed a modest 2-fold increase with 10% DGS-NTA (Ni^2+^)-containing liposomes (Fig. 2A). Drp1 functions in organelle division and we recently recreated a Drp1-catalyzed membrane fission reaction on supported membrane nanotubes (SMrT) (Kamerkar et al. 2018a). Fission requires robust stimulation of Drp1’s GTPase activity. Consistent with results from GTPase assays (Fig. 2A), Drp1 caused dramatic fission of SMrT templates with 30% CL but not with templates containing 10% DGS-NTA(Ni^2+^) (Fig. 2B,C). These results signify that membrane recruitment via DGS-NTA(Ni^2+^) does not mimic native Drp1-lipid interactions that allow it to function in membrane fission.

**Fig. 2.**
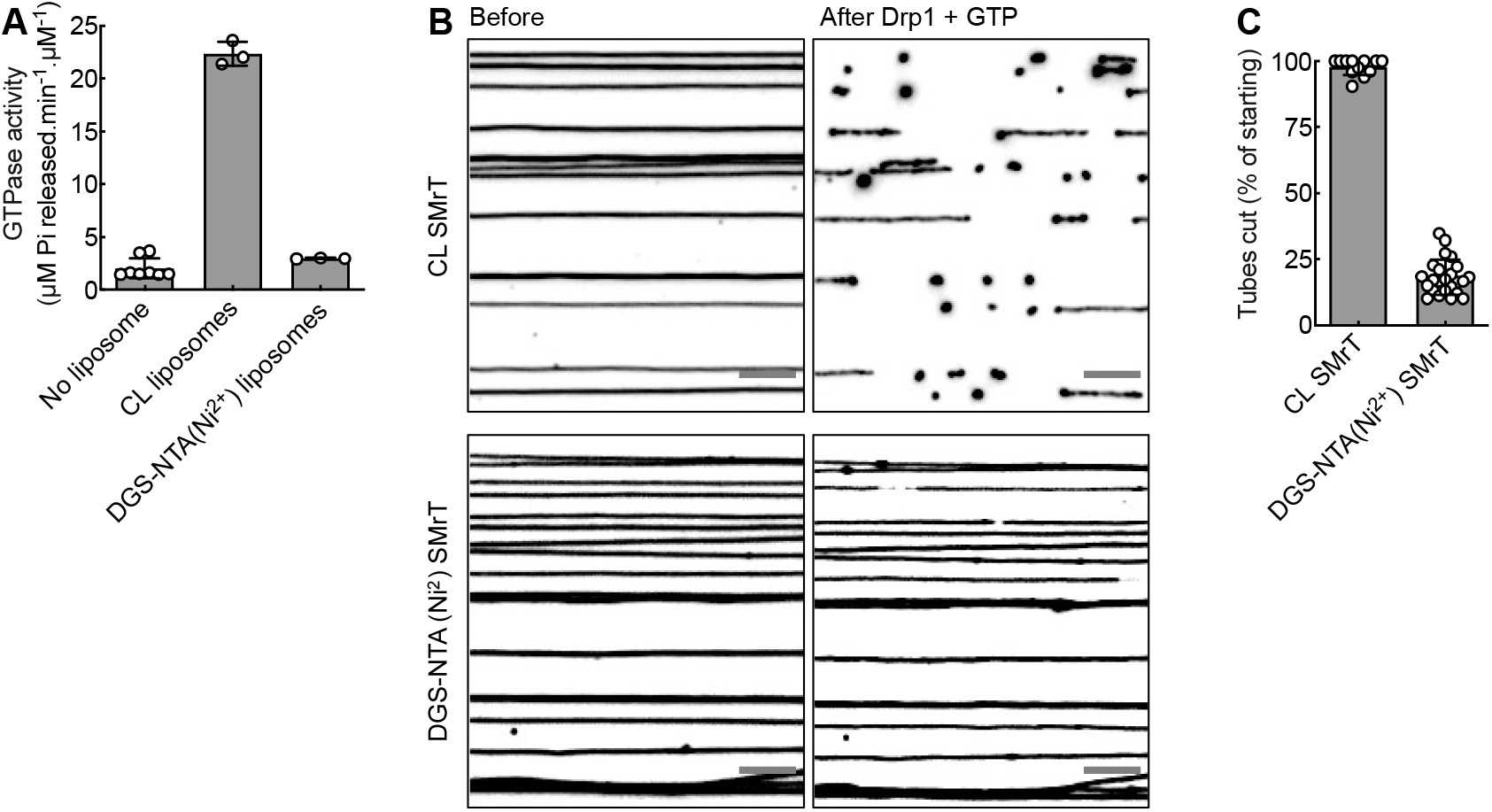
DGS-NTA(Ni^2+^)-recruited Drp1 is functionally inactive. (A) Drp1-catalyzed GTP hydrolysis in the absence and presence of 30 mol% CL- and 10 mol% DGS-NTA(Ni2+)-containing liposomes. Data represent the mean ± SD of at least three independent experiments. (B) Representative fluorescence micrographs of supported membrane nanotubes (SMrT) containing 30 mol% CL (top) and 10 mol% DGS-NTA(Ni2+) (bottom) imaged before and after addition of Drp1 with GTP. Fission is seen as severing of the nanotubes. Images are inverted in contrast and scale bars = 10 μm. (C) Quantitation of Drp1-catalyzed membrane fission of SMrTs containing 30 mol% CL and 10 mol% DGS-NTA(Ni^2+^). Data represents the mean ± SD of the percentage of severed tubes in a microscope field averaged across multiple fields from two independent experiments.

Proteins that bind NTA(Ni^2+^) generally do so through a contiguous stretch of metal-binding residues (Bolanos-Garcia and Davies 2006; Sudan et al. 2015; Yang et al. 2019). In the case of 6xHis tags interacting with NTA(Ni^2+^), the NTA moiety occupies four of the six coordination sites on the metal ion while the remaining two are occupied by any two of the six imidazole moieties in the His tag (Hochuli and Piesecki 1992; Zhao et al. 2010). To understand how Drp1 binds DGS-NTA(Ni^2+^), we systematically tested conditions that compete with or perturb interactions between histidine residues and NTA(Ni^2+^). PLiMAP experiments where 10 mol% DGS-NTA(Ni^2+^)-containing liposomes were incubated with Drp1 in presence of increasing concentrations of a random 6xHis tagged protein (6xHis-mCherry) showed a decrease in Drp1 fluorescence and an increase in 6xHis-mCherry fluorescence, implying that a 6xHis tag competes with Drp1 binding to DGS-NTA(Ni^2+^) (Fig. 3A,B). Interestingly, Drp1’s binding to DGS-NTA(Ni^2+^) could not be competed out entirely and a significant fraction remained membrane-bound even in presence of 10-fold molar excess of the 6xHis mCherry. This has implications on the use of DGS-NTA(Ni^2+^) in reconstitution experiments with 6xHis-tagged constructs as very likely there will persist a Drp1 population that is directly engaged with DGS-NTA(Ni^2+^). Drp1’s binding to DGS-NTA(Ni^2+^) was also reduced in presence of increasing concentrations of the histidine side chain moiety Imidazole (Fig. 3C,D), indicating that this interaction is mediated by histidine residues on the protein forming a coordination complex with the Ni^2+^ ion (Fig. 3C,D). Lastly, striping-off the Ni^2+^ ion on DGS-NTA with EDTA abolished binding (Fig. 3C,D), implying that Drp1 binds the Ni^2+^ ion and not the chelator lipid DGS-NTA. Together, these results indicate that Drp1 is a metal-binding protein and that such interactions are mediated via histidine residues in the protein.

**Fig. 3.**
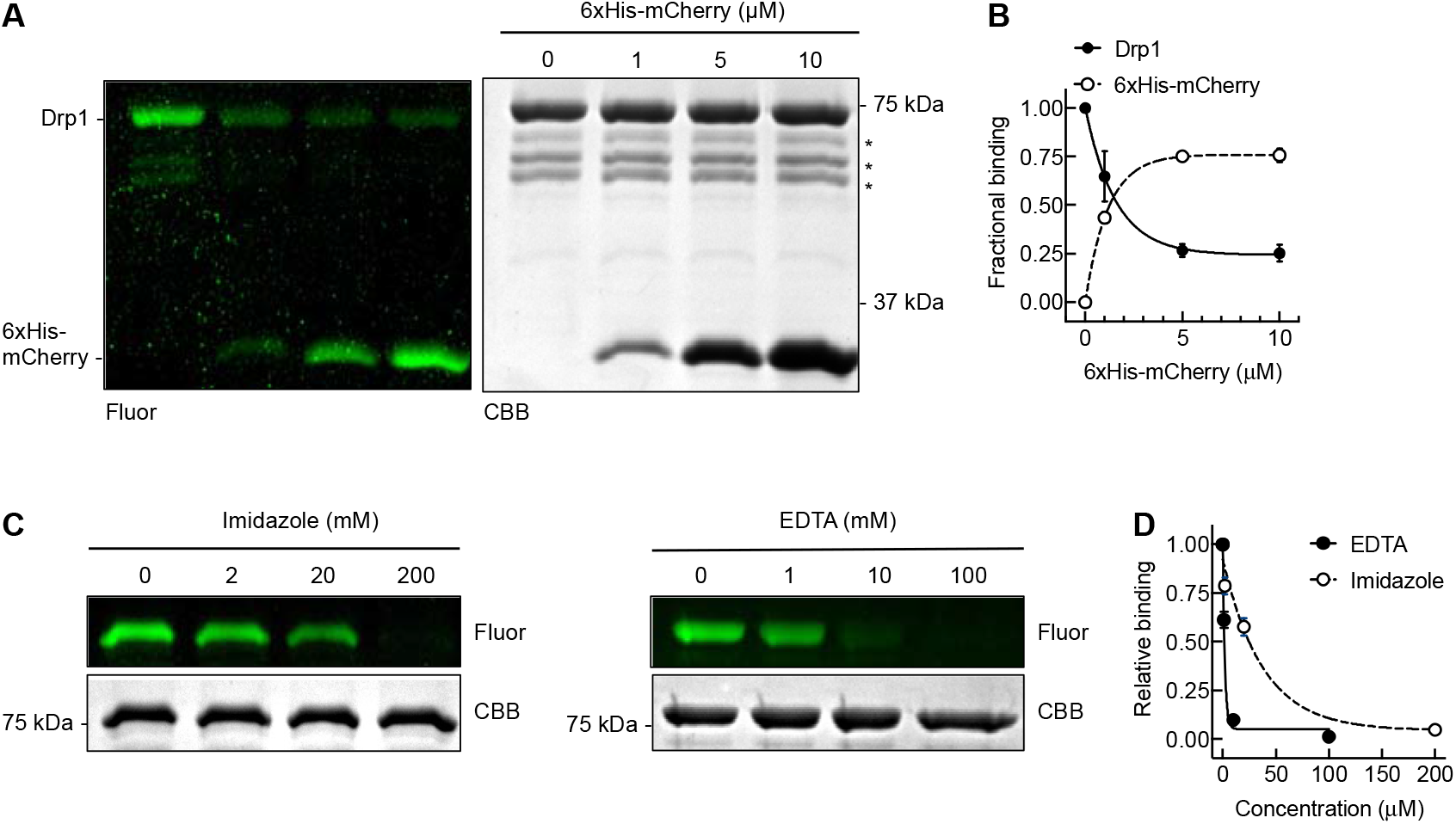
Determinants of Drp1-DGS-NTA(Ni^2+^) interactions. (A) Results from a representative PLiMAP experiment showing in-gel fluorescence (Fluor) and Coomassie brilliant blue (CBB) staining of Drp1 and 6xHis mCherry with 10 mol% DGS-NTA(Ni^2+^)-containing liposomes. Asterix mark degraded Drp1. (B) Quantitation of ingel fluorescence of Drp1 and 6xHis mCherry. Data is represented as fractional liposome binding, defined as the ratio of Drp1 or 6xHis-mCherry fluorescence to the sum of Drp1 and 6xHis-mCherry fluorescence. Data represents mean ± SD of three independent experiments. (C) Results from a representative PLiMAP experiment showing in-gel fluorescence (Fluor) and Coomassie brilliant blue (CBB) staining of Drp1 in presence of 10 mol% DGS-NTA(Ni^2+^) liposomes with increasing concentrations of Imidazole and EDTA. (D) Quantitation of in-gel fluorescence of Drp1 in presence of imidazole and EDTA. Data represents the mean ± SD of three independent experiments.

Next, we tested if Drp1’s propensity to bind metals chelated on a surface is apparent in a native context. For this, we incubated brain lysates with increasing amounts of NTA(Ni^2+^) chelator beads, sedimented them and blotted the supernatant for Drp1. As seen in Fig. 4A, brain lysates incubated with the NTA(Ni^2+^) beads (but not with NTA beads alone) showed a dose-dependent reduction in Drp1 levels in the supernatant, implying efficient pull down of Drp1. Consistently, Drp1 was found bound to NTA(Ni^2+^) beads but not to NTA beads (Fig. 4B).

**Fig. 4.**
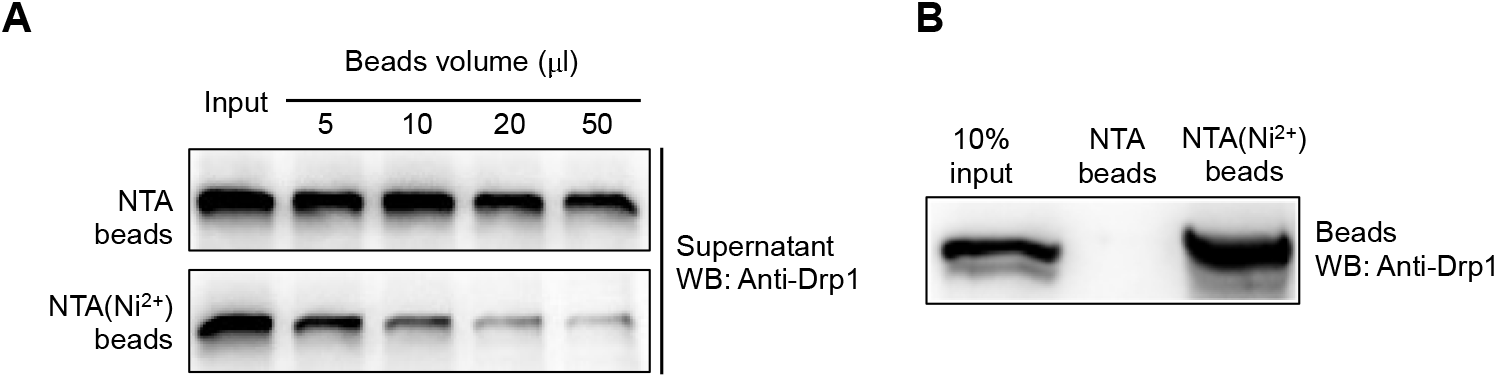
Native Drp1 interacts with Ni^2+^. (A) Representative western blots of Drp1 in the supernatant of a reaction where brain lysate was incubated with increasing amounts of NTA beads or beads charged with Ni^2+^. (B) Representative western blot of Drp1 pulled down with 50 μl of NTA beads or beads charged with Ni^2+^.

Our results show that membrane recruitment via DGS-NTA(Ni^2+^) does not mimic native Drp1-lipid interactions that allow it to function in membrane fission (Fig. 2). This could either be an outcome of constrained sub-optimal orientation of Drp1 on the membrane or that metal binding inhibits Drp1 functions, or both. To directly test if metal binding affects Drp1 functions, we tested GTPase and membrane fission activities on CL-containing templates in presence of free Ni^2+^. Remarkably, Drp1’s GTPase activity with CL liposomes showed a dramatic reduction with increasing concentrations of Ni^2+^ added to the buffer (Fig. 5A). The stimulated GTPase activity with Ni^2+^ (1 mM) in solution reduced to basal levels seen with Drp1 in the absence of liposome. On the other hand, the basal GTPase remained largely unaffected (Fig. 5A). Consistently, presence of Ni^2+^ (1 mM) abolished Drp1’s fission activity on 30% CL-containing SMrT templates (Figs. 5B,C). Together, these results indicate that the metal significantly inhibits Drp1 functions.

**Fig. 5.**
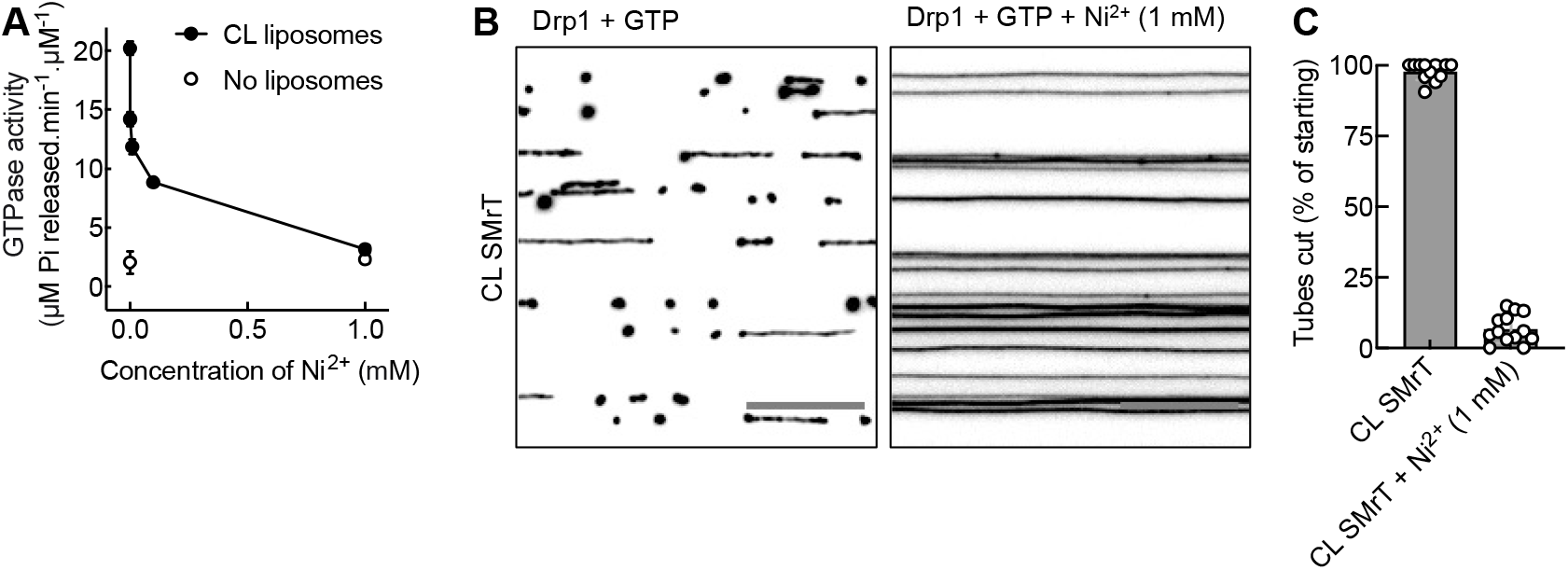
Ni^2+^ inactivates Drp1. (A) Drp1-catalyzed GTP hydrolysis in the absence and presence of increasing concentrations of Ni^2+^ assayed with and without CL liposomes. Data represent the mean ± SD of at least two independent experiments. (B) Representative fluorescent micrographs of SMrT containing 30 mol% CL after addition of Drp1 (1 μM) with GTP (1 mM) in the absence (left) or presence (right) of Ni^2+^ (1 mM). Images are inverted in contrast and scale bar = 10 μm. (C) Quantitation of Drp1-catalyzed membrane fission in the absence and presence of Ni^2+^ (1 mM) on 30 mol% CL SMrTs. Data represents the mean ± SD of the percentage of severed tubes in a microscope field averaged across multiple fields from two independent experiments.

## Discussion

In this report, we characterize an intrinsic metal ion-binding propensity in both native and recombinant Drp1. This property can be exploited to deplete or purify Drp1 from native tissues using standard resins used in immobilized metal affinity chromatography (IMAC). It is possible that the buffers used in lysate preparation could have removed any bound metal ion thus explaining why native Drp1 showed no binding to the IMAC resin devoid of metals. Drp1 is expressed as multiple splice variants in a tissue-specific manner (Itoh et al. 2018). Current strategies of correlating functions of these spice variants to tissue physiology rely on RT-PCR based identification of the splice variants in specific tissues. The present work utilizes the shortest splice variant (isoform 3) indicating that metal binding should be a property conserved among all splice variants. An IMAC-based pulldown and subsequent mass spectrometry could therefore represent a facile approach to identify tissuespecific expression of the splice variants and their abundance.

While we began this work with the intention of confirming an earlier reported interaction between Drp1 and metal-bound chelator lipid, we find that free metal has significant consequences on Drp1 functions. The presence of Ni^2+^ inactivates Drp1’s stimulated GTPase activity and renders it deficient in membrane fission. Liposome-stimulated GTPase activity is attributed to self-assembly of the protein on the membrane surface and concomitant reorientation of catalytic resides in the G-domain. Since we find that the metal selectively reduces stimulated but not basal GTPase activity, it is conceivable that the metal binding site is located on a self-assembly interface of the protein. Future work will expand the scope of the present investigation by testing binding specificity with physiologically relevant heavy metals and identification of the metal-binding site(s). Such work could have wider implications as heavy metals are known to affect mitochondrial structure and function (Dineley et al. 2003; Zischka and Borchard 2019). An exciting possibility is that such effects are mediated in part by the metal binding to and inhibiting Drp1 function.

## Acknowledgements

We thank Gregor Jose for preliminary work on testing the binding of Drp1 to DGS-NTA(Ni^2+^). We thank Pucadyil Lab members for valuable comments on the manuscript.

## Author Contributions

KR and TJP conceptualized the study. KR performed all experiments. KR and TJP analyzed data and prepared the manuscript.

## Declarations

Conflict of interest: The authors declare no conflict of interest.

